# Enhancement strategies for the microbial protein production of nitrogen-fixing hydrogen-oxidizing bacterial community

**DOI:** 10.1101/2024.08.07.607053

**Authors:** Haoran Wang, Daping Li

## Abstract

The feasibility and superiority of utilizing nitrogen-fixing hydrogen-oxidizing bacteria (NF-HOB) for microbial protein (MP) production have been proposed. This study simulated air as the source of nitrogen and oxygen and enhanced production efficiency by employing key strategies, including continuous gas supply, selecting a higher hydrogen-to-oxygen ratio, designing initial community structures and domestication, and exploring appropriate hydraulic retention times (HRT) in continuous culture. The results demonstrated that sequencing batch culture achieved an MP titer of 0.98 g/L, with N_2_ fixation efficiency surpassing natural nodules by two orders of magnitude at 1.6 mg N_2_ per g dry biomass per hour. Under an HRT of 48 hours, MP volumetric productivity reached 2.297 mg/(L·h), accompanied by a maximum biomass yield of 0.11 g CDW/g COD-H_2_. The high abundance of nitrogenase may provide crucial functional support, enabling the NF-HOB community to exhibit potential beyond previous understanding.

## Introduction

Hydrogen-oxidizing bacteria (HOB) have recently attracted significant attention for their ability to utilize renewable hydrogen energy for carbon dioxide fixation and their potential in the synthesizing high-quality microbial protein (MP) (<u>Matassa et al.,</u> <u>2015</u>; <u>Woern and Grossmann, 2023</u>). However, a majority of HOB employed for MP production still necessitate energy-intensive inputs (<u>Hu et al., 2020</u>; <u>Sherbo et al.,</u> <u>2022</u>), particularly ammonium salts derived from the Haber-Bosch process as an optimal nitrogen source. This practice leads to carbon dioxide emissions, contradicting the inherent advantages of autotrophic bacteria as nutrient producers.

Therefore, the advantages of nitrogen-fixing hydrogen-oxidizing bacteria (NF-HOB), also known as “H-C-N microorganisms” (<u>Sherbo et al., 2022</u>), have been further proposed. This group includes members from the following genera: *Azospirillum* (*Azospirillum lipoferum*), *Ancylobacter* (*Ancylobacter aquaticus*, *Ancylobacter vacuolatus*), *Xanthobacter* (*Xanthobacter autotrophicus*, *Xanthobacter flavus*, *Xanthobacter variabilis*), *Bradyrhizobium* (*Bradyrhizobium japonicum*), *Azohydromonas* (*Azohydromonas lata*), and *Derxia* (*Derxia gummosa*) (<u>Hu et al., 2022</u>; <u>Lu et al., 2020</u>; <u>Malik and Schlegel, 1981</u>; <u>Scott et al., 2021</u>). Cultivation experiments are essential for identifying NF-HOB, as they allow for determining their ability to simultaneously fix CO_2_ and N_2_ under hydrogen autotrophic conditions for growth. Utilizing NF-HOB for MP production represents a highly sustainable approach that effectively harnesses the abundant atmospheric resources of N_2_ and O_2_, integrating hydrogen production technology with MP synthesis regardless of climate or geographical constraints. Notably, when solar energy is used for hydrogen production technology, the solar-to-biomass energy conversion efficiency of NF-HOB was more than double that observed in symbiotic N_2_-fixing soybeans (<u>Hu et al., 2020</u>).

Currently, cultivating NF-HOB is still in the experimental phase within small-scale systems. Hu et al. successfully achieved enrichment of NF-HOB community under headspace gas supply conditions (2% O_2_, 10% H_2_, 10% CO_2_, 78% N_2_) in serum bottles, resulting in a protein-rich biomass with an amino acid profile comparable to that of MP derived from ammonium-based HOB. However, biomass growth under this system was significantly constrained, with the titer of nitrogen content in biomass being less than 12 mg/L (<u>Hu et al., 2020</u>). Recently, a comparative experiment was conducted in a continuous gas supply system to evaluate nitrogen source supply conditions. The results showed a slightly higher N_2_ fixation amount in the simple device compared to previous studies, confirming the potential of continuous feed gas for enriching NF-HOB. Importantly, no significant difference was observed in the absolute abundance of hydrogenase or Rubisco between the NF-HOB community and the control community supplied with ammonia nitrogen, suggesting that the synthesis of essential functional enzymes was not limited. Furthermore, it is noteworthy that air can potentially substitute separately supplied N_2_ and O_2_ based on a feed gas composition of H_2_/O_2_/CO_2_/N_2_ = 6/1/1.56/4.28 (v/v/v/v) (<u>Wang et al., 2024</u>).

It should be emphasized that, considering the future potential of large-scale utilization of NF-HOB for MP production, developing a functionally stable microbial community offers greater advantages in mitigating external pollution compared to pure strains (<u>Matassa et al., 2015</u>). Optimizing the hydrogen-oxygen ratio (R_H/O_) in the feed gas and exploring cultivation in a continuous reactor are promising strategies for enhancing MP production based on NF-HOB community, considering the influence of feed gas component proportions on functional enzyme activity and the promoting effect of continuous culture mode for the ammonium-based HOB community (<u>Matassa et al., 2016</u>).

This study employed a more efficient gas-liquid mass transfer fermentation system and incorporated a N_2_-O_2_ mixed gas with a ratio of 4:1 to simulate the feasibility of air inclusion in the feed gas. Furthermore, deliberate design and acclimation were applied to the initial bacterial community structure to enhance its N_2_ fixation activity. Subsequently, the performance of NF-HOB community in producing MP under slightly lower or higher R_H/O_ (3.5 and 7, respectively) compared to previous studies (R_H/O_ = 5 or 6) was investigated in sequencing batch culture, with ammonium provision as the comparative condition. Additionally, the effective hydraulic retention time (HRT) was investigated in continuous cultivation while simultaneously assessing volumetric productivity and biomass yield. An analysis of the impact of these enhancement strategies on the microbial community structure was conducted, accompanied by a description of functional abundance characteristics to provide mechanistic support. The research findings revealed significant potential for synthesizing MP based on NF-HOB, surpassing previous understanding.

## Material and methods

### Design and domestication of initial microbial community

The NF-HOB community obtained from the previous study was combined with the laboratory-preserved NF-HOB strains (*Xanthobacter flavus* strain CIB1 [GenBank: OP108817.1] and unclassified Xanthobacteraceae bacterium strain CIB2) and subsequently inoculated into a device previously used for enriching NF-HOB (<u>Wang et</u> <u>al., 2024</u>). The flow rates of each component in the continuously supplied feed gas were: H_2_ 81 mL/min, O_2_ 13.5 mL/min, CO_2_ 21 mL/min, N_2_ 57.75 mL/min (R_H/O_ = 6).

The components of the base medium included: K_2_HPO_4_ 1 g/L, KH_2_PO_4_ 0.5 g/L, NaHCO_3_ 2 g/L, MgCl_2_·6H_2_O 0.12 g/L, CaCl_2_ 0.0258 g/L, NaVO_3_·H_2_O 50 mg/L, Na_2_MoO_4_·2H_2_O 25 mg/L, FeSO_4_·7H_2_O 50mg /L, vitamin solution and trace element solution both added at 1mL/L. Cultivation was carried out at a constant temperature of 30 for one month.

The cultivation process demonstrated a significant increase in turbidity, indicating the community’s potential for growth under hydrogen autotrophy and nitrogen fixation conditions after domestication, making it suitable for subsequent quantitative experiments.

### Sequencing batch culture

The gas fermentation system constituted the primary component of the reactors (see supplementary material). Each glass fermentation tank was equipped with three agitators, each featuring four-blade impellers, as well as a thermometer, a pH meter, and an aeration device. The stirring speed and temperature were controlled by the fermentation system at 500 rpm and 30 , respectively. The gas cylinder continuously supplied H_2_, N_2_-O_2_ mixed gas (4:1, simulated air), and CO_2_, which were thoroughly mixed after passing through pipelines, flow-stabilizing valves, and rotameters in sequential order as previously described (Wang et al., 2024). The mixture was then introduced into the fermentation tank. The flow rate of each gas component was adjusted by manipulating the valve and rotameter to precisely regulate R_H/O_. Exhaust gases were collected using gas collection bags that demonstrated excellent airtightness.

In order to better evaluate the growth trend, two fermentation tanks were utilized with the base medium and base medium supplemented with 2 g/L NH_4_Cl (2 L, pH adjusted to 7.5 using 1 mol/L hydrochloric acid prior to inoculation). Consequently, ammonium assimilation conditions were employed as the control for nitrogen fixation conditions. Consistent initial inoculation and gas intake were maintained in both tanks.

Under low R_H/O_ (3.5) conditions, the total flow rates of H_2_, “air”, and CO_2_ prior to distribution into the two fermentation tanks were 42 mL/min, 60 mL/min, and 20 mL/min, respectively. In contrast, under high R_H/O_ (7) conditions, the flow rate of “air” was adjusted to 30 mL/min.

During the batch cultivation period, daily samples were collected from the fermentation tank to quantify total chemical oxygen demand (COD_t_) and cell dry weight (CDW) for biomass titer characterization, while monitoring pH. The collected samples were filtered using a membrane filter (0.22 μm Nylon66, JINTENG, China) and then analyzed for soluble chemical oxygen demand (COD_s_), nitrate nitrogen, nitrite nitrogen, and ammonia nitrogen. Biomass measurement at the end of cultivation necessitated transferring the bacteria that had been suspended and adhered to the tank wall due to agitation into the liquid phase and thoroughly homogenizing before sampling.

### Continuous culture

For the continuous reactor, the liquid flow rate was controlled through the control panel (see supplementary material), allowing automated operation of the peristaltic pumps and ensuring equal amounts of fresh base medium and culture solution in and out of the system. The liquid phase volume was stabilized at 2 L; thus, the required liquid flow rate can be determined based on HRT. The other cultivation conditions were consistent with those used in batch culture.

The community that exhibited stable proliferation in the sequencing batch culture was utilized as the inoculum source. Samples were collected from both the inner liquid phase and the collector for liquid discharge in order to measure COD_t_ (inner) and daily average COD_t_ (outlet), respectively.

### Quantification of H_2_ utilization

Referring to previous NF-HOB research, the quantification of H_2_ consumption corresponding to biomass growth was conducted through cultivation experiments in anaerobic flasks to determine the biomass yield (<u>Hu et al., 2020</u>; <u>Wang et al., 2024</u>). The proportion of each component in the headspace gas followed that of the continuous gas supply system and was cultured in a constant temperature shaker (30 , 180 rpm). The headspace gas was updated every two days with the total effective cultivation time of 6 days. For NF-HOB communities obtained from sequencing batch culture and continuous culture, measurements were taken for CDW, COD_t_, and pH at both the beginning and end of the culture period. This experiment was performed in triplicate.

### Analysis and characterization of MP

The carbon and nitrogen content in the biomass was determined using an elemental analyzer (vario EL cube, ELEMENTAR, Germany). The obtained nitrogen content was then multiplied by a conversion coefficient of 6.25 to determine the MP content (<u>Matassa et al., 2016</u>). The dietary amino acid composition of the biomass was determined by Baihui Biotech (Chengdu, China).

### Microbial community analysis

The bacterial solutions to be analyzed were each sampled with a minimum volume of 50 mL, followed by centrifugation for supernatant removal and subsequent storage at -80 °C. DNA extraction of culture samples and Illumina NovaSeq sequencing were performed by Novogene (Beijing, China). MetaStat analysis was used to identify species with significantly different abundances between groups. Functional annotation of PICRUST2 was employed to reflect the abundance of important functional enzymes or their coding genes.

### Analytical methods and calculations

The concentrations of COD_t_ and COD_s_ were determined using the COD detection kit (Titric, China). The pH value was determined using a pH meter (Seven2Go, METTLER TOLEDO, Switzerland). The concentrations of NH^+^_4_ -N, NO_3_^-^ -N, and NO^-^_2_ -N were determined using Nessler reagent spectrophotometry, ultraviolet spectrophotometry, and N-(1-naphthyl)-ethylenediamine photometry techniques, respectively. CDW determination was accomplished by desiccating and quantifying a fixed volume sample, as previously described (<u>Wang et al., 2024</u>).

In the liquid phase, the total nitrogen (TN) is calculated as: TN = Total nitrogen content of biomass (TN_biomass_) + NH_4_^+^-N (AN) + NO_3_^-^-N + NO_2_^-^-N; soluble total nitrogen (TN_s_) = AN + NO_3_^-^-N + NO_2_^-^-N, TN_biomass_ (mg/L) = N content (%) ^×^ CDW (mg/L).

NH ^+^_4_ -N conversion efficiency (%) = (initial AN concentration – final AN concentration) / initial AN concentration; NH^+^_4_ -N utilization efficiency (%) = increased TN_biomass_ / (initial AN concentration – final AN concentration). Biomass yield (g CDW/g COD-H_2_) = increased CDW (mg) / [8 (g COD/g H_2_) × reduced H_2_ (mg)].

## Results and discussion

### A high hydrogen-oxygen ratio facilitated microbial protein production with high titers

O_2_ exerts inhibitory effects on both hydrogenase (<u>Greening et al., 2023</u>; <u>Zacarias</u> <u>et al., 2019</u>) and nitrogenase (<u>Allen et al., 2019</u>), while also competing to inhibit the CO_2_ reduction activity of Rubisco (<u>Poudel et al., 2020</u>). However, low concentrations of O_2_ can result in insufficient electron acceptors. Therefore, optimizing the R_H/O_ of feed gas is a primary strategy to consider.

The autotrophic growth of *Xanthobacter* in a gas mixture with an H_2_/O_2_/CO_2_ ratio of 7/2/1 was initially documented (<u>Wiegel et al., 1978</u>). Additionally, the H/O ratio observed in the biomass composition of various HOB strains ranged from 2.71 to 3.77 (<u>Ishizaki and Tanaka, 1990</u>; <u>Yu and Lu., 2019</u>; <u>Zhang et al., 2020b</u>). However, results from sequencing batch culture experiments revealed that an R_H/O_ of 3.5 solely facilitated ammonium assimilation without supporting biomass growth during N_2_ fixation.

Under N_2_ fixation conditions, COD_t_ gradually increased, reaching only 158 mg/L over 10 days, while pH fluctuated around 7.6. In the presence of ammonia nitrogen, both COD_t_ and CDW rapidly increased, reaching 4552 mg/L and 2360 mg/L, respectively, on the 10th day. Notably, efficient CO_2_ absorption during this period caused the culture medium’s pH to decline from 7.88 to 6.12 (Fig. 1A). The observed difference surpassed previous studies’ outcomes, likely due to the lower R_H/O_.

**Fig. 1.**
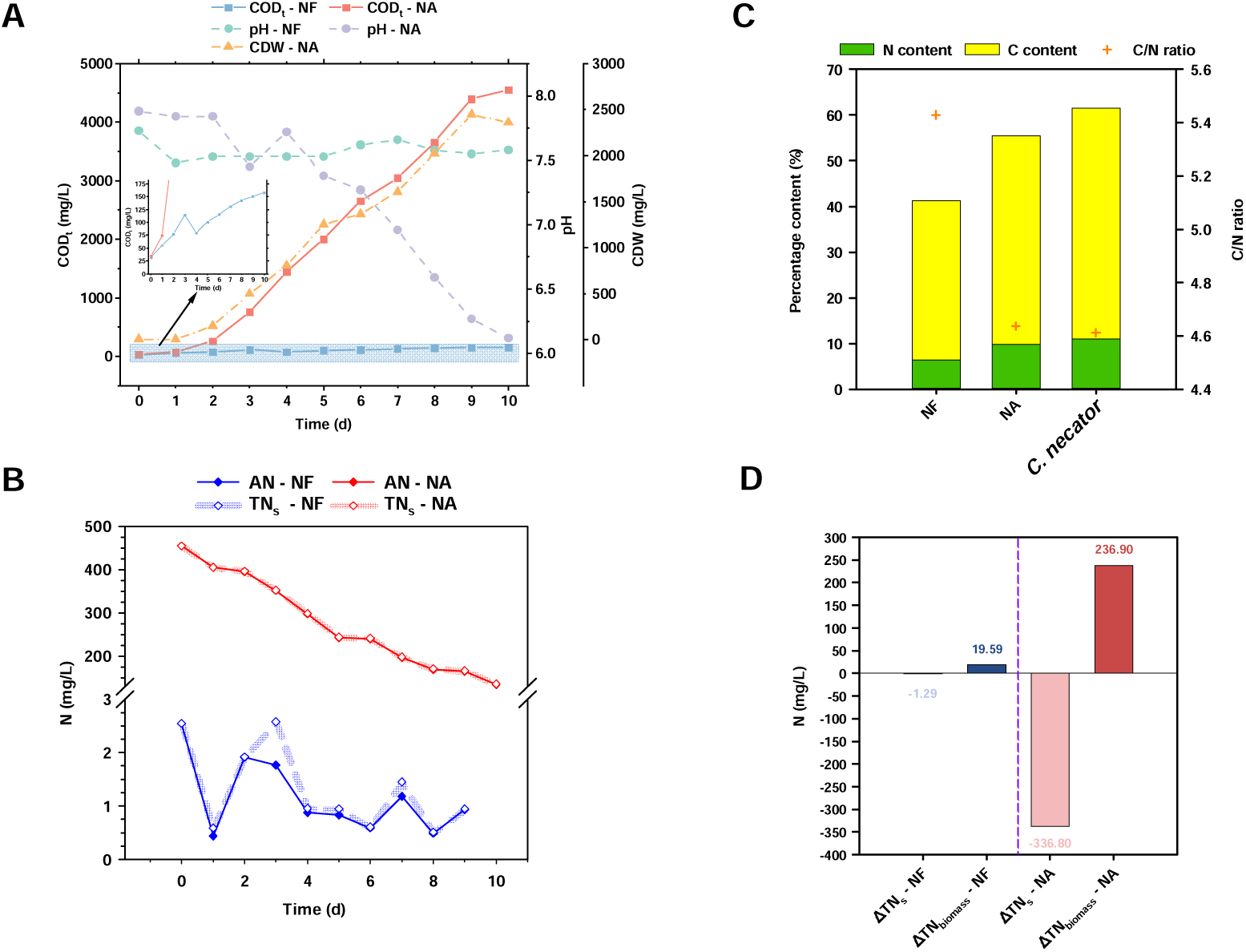
Characterization of biomass growth, pH change, and N element flow during sequencing batch culture of HOB communities (NF: NF-HOB community; NA: ammonium-assimilation HOB community) at R_H/O_ = 3.5: (A) COD_t_ concentration, pH change, and CDW concentration; (B) NH^+^_4_ -N (AN) and TN_s_ concentrations; (C) carbon and nitrogen content and C/N ratio in biomass (*C. necator* as reference); (D) changes in TN_s_ and TN_biomass_.

The nitrogen transformation process aligned well with expectations, evidenced by the low concentration of TN_s_ (mainly NH^+^_4_ -N) in the medium under N_2_ fixation conditions (Fig. 1B) and a biomass titer of 305.56 mg/L (after three weeks of cultivation) with a nitrogen content of 6.41% (Fig. 1C). Consequently, most reduced N_2_ was assimilated into the biomass, leading to a significant increase of 19.59 mg/L TN_biomass_ (^Δ^TN_biomass_ – NF shown in Fig. 1D), surpassing previous research findings. The ammonium assimilation control exhibited remarkable efficiency in absorbing ammonium, achieving a conversion efficiency of 70.3% over ten days while maintaining very low levels of nitrate or nitrite nitrogen (Fig. 1B). Finally, the residual NH^+^_4_ -N was measured at 116.5 mg/L, and CDW displayed a titer of 2410 mg/L (pH = 5.29) with a nitrogen content of 9.83% (Fig. 1C), resulting in an impressive ammonium utilization efficiency of 70.1% (Fig. 1D). Furthermore, biomass obtained under ammonium assimilation conditions exhibited a carbon and nitrogen composition more similar to that of *Cupriavidus necator* (previously named as *Alcaligenes europhus*) strain ATCC 17697T (<u>Ishizaki and Tanaka, 1990</u>), widely used as a reference for elucidating the stoichiometric relationship of HOB, rather than under N_2_ fixation conditions (Fig. 1C).

Adjusting R_H/O_ to 7 significantly enhanced the N_2_-fixing capacity of the NF-HOB community, leading to increased biomass accumulation and MP synthesis. In the first batch, following a lag period of approximately 9 days, the NF-HOB community exhibited rapid linear biomass growth, accompanied by a significant decrease in pH (Fig. 2A). By the 15th day, COD_t_ had increased impressively by 426% to 1299 mg/L. TN_s_ were remarkably low, primarily composed of ammonia nitrogen and nitrate nitrogen, as shown in Fig. 2D. The COD_t_ and CDW of the ammonium-assimilation HOB community continued to increase steadily; however, at a significantly slower rate compared to low R_H/O_ conditions, reaching values of 2342 mg/L and 1760 mg/L on the 10th day, respectively. Consequently, there was also a reduction in pH decline (Fig. 2A). Surprisingly, the concentration of NH^+^_4_ -N or TN_s_ increased by 82.5 mg/L in the initial 3 days (Fig. 2D), while biomass exhibited growth, suggesting this was due to N_2_ fixation rather than bacterial lysis. This phenomenon is similar to observations in previous studies (<u>Wang et al., 2024</u>). The biomass accumulation of the NF-HOB community at the end of this batch was significantly enhanced, CDW reaching 1088.3 mg/L, thereby substantially reducing the disparity in biomass accumulation between NF-HOB and ammonium-assimilation HOB.

**Fig. 2.**
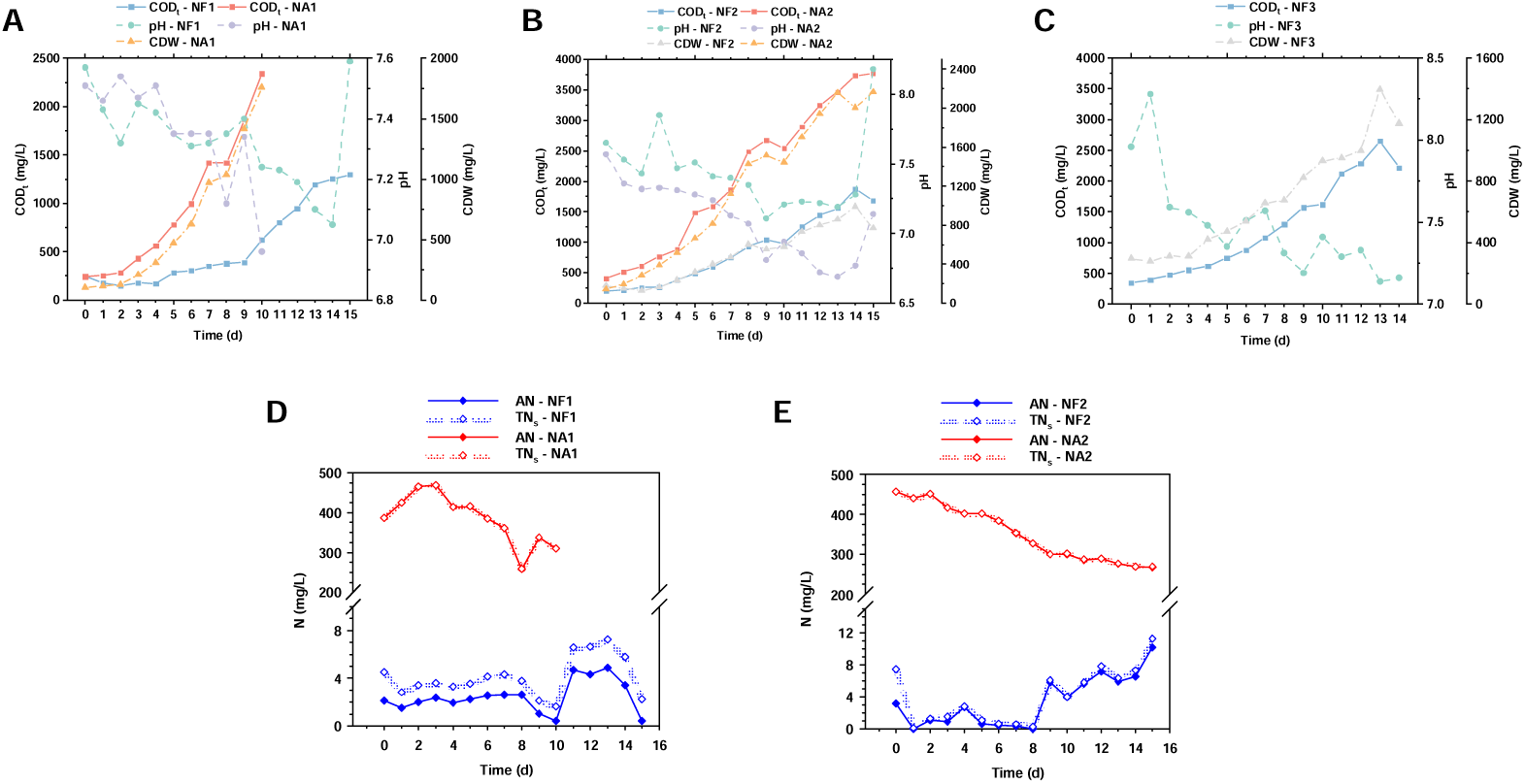
Characterization of biomass growth, pH change, and soluble-nitrogen variation during sequencing batch culture of HOB communities (NF: NF-HOB community; NA: ammonium-assimilation HOB community) at R_H/O_ = 7: COD_t_ concentration, pH change, and CDW concentration in the first (A), second (B) and third (C) batches; NH^+^_4_ -N (AN) and TN_s_ concentrations in the first (D) and second (E) batches.

As for the second batch, over 14 days the NF-HOB community demonstrated remarkable increases in COD_t_ by 815% and CDW by 450% (Fig. 2B). During the latter half of the cultivation period, both NH^+^_4_ -N and TN_s_ concentrations fluctuated and then increased, surpassing 10 mg/L on the 15th day (Fig. 2E). The growth rate of COD_t_ and CDW in the ammonium-assimilation HOB community closely approximated that observed in the first batch, while also showing fluctuations in NH^+^_4_ -N concentration.

Excitingly, after extending the culture for an additional 5 days, NF-HOB achieved a CDW titer of 1713 mg/L upon collection of adherent cells and bacterial fluid, closely approaching the final titer of ammonium-assimilation HOB at 1725 mg/L – it is not uncommon for biomass to decrease with prolonged cultivation.

The biomass growth trend of the NF-HOB community in the third cultivation batch was similar to that observed in the second batch, demonstrating even greater increments in COD_t_ and CDW (Fig. 2C). The final concentration of COD_t_ was 3312 mg/L, with 2880 mg/L from COD_biomass_ (the proportion of COD_biomass_, always the main component, tended to be highest during the middle stage of cultivation, see supplementary material), resulting in a CDW titer of 2182 mg/L.

Elemental analysis revealed that the nitrogen content of NF-HOB biomass was slightly lower than that of ammonium-assimilation HOB, resulting in a higher C/N ratio. MP content significantly increased compared to results obtained under low R_H/O_, ranging from 45.06% to 53.88% (Fig. 3A). By quantifying the nitrogen content in each component, it can be inferred that the NF-HOB community fixed more than 278 mg of N_2_ in both of the last two batches (Fig. 3B), demonstrating substrate-based N_2_ fixation efficiencies of 1.6 and 1.2 mg N_2_ per g dry biomass per hour, respectively. This level exceeds that of natural nodules by two orders of magnitude (<u>Lu et al., 2020</u>). Fig. 3B also illustrates variations in TN_s_ and changes in TN_biomass_ for ammonium-assimilation HOB communities, further substantiating the occurrence of N_2_ fixation under conditions characterized by high R_H/O_, which induced fluctuations in TN_s_ levels (Fig. 2D and 2E).

**Fig. 3.**
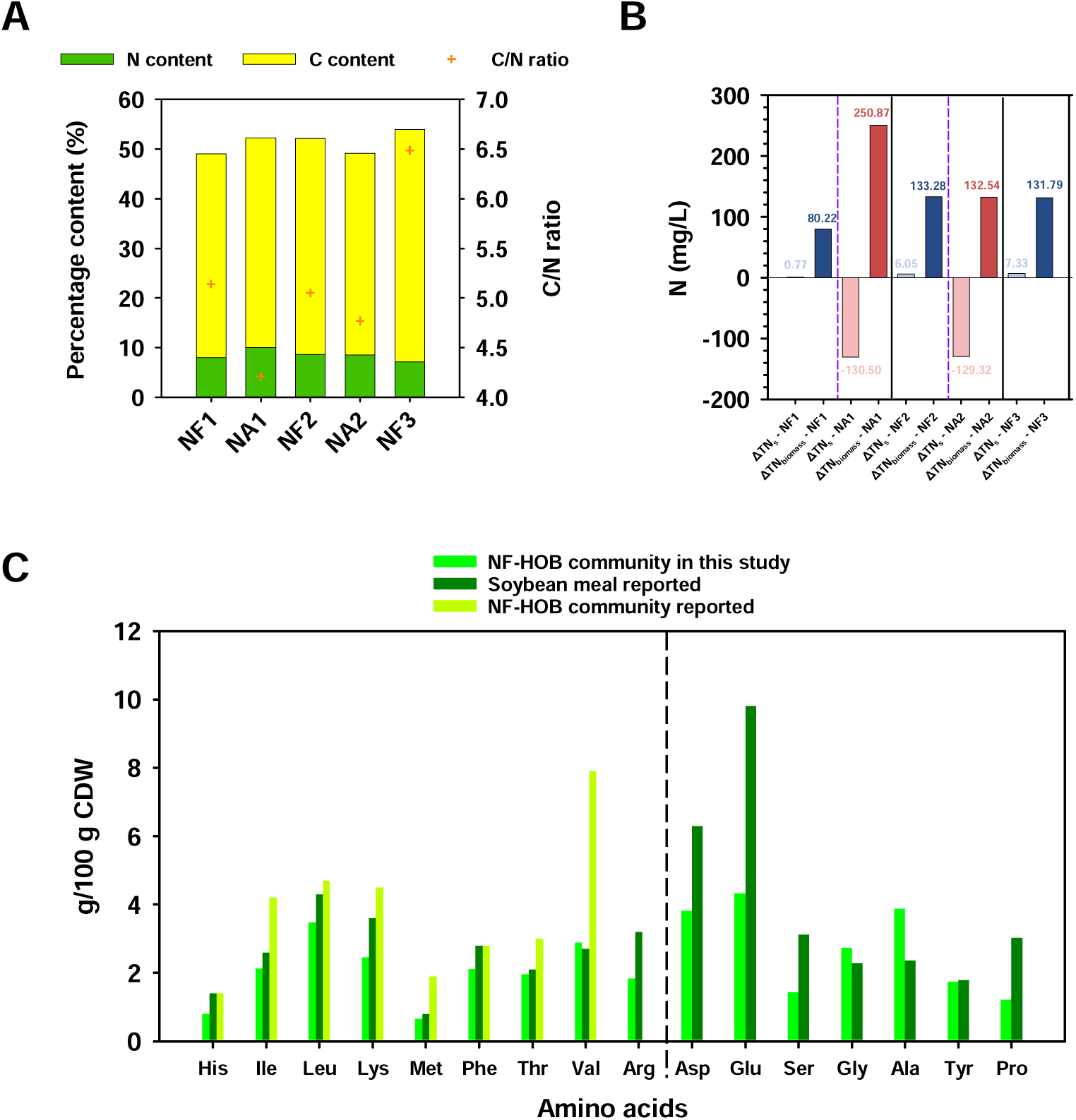
(A) Carbon and nitrogen content and C/N ratio in biomass of HOB communities at R_H/O_ = 7 and (B) changes in TN_s_ and TN_biomass_ during each sequencing batch; NF1, NF2, and NF3 represent the NF-HOB community in the first, second, and third batches, respectively; NA1 and NA2 represent the ammonium-assimilation HOB community in the first and second batches, respectively. (C) Amino acids profile of high-titer biomass (this study), compared with soybean meal reported and the NF-HOB community previously enriched within small-scale systems (His: Histidine; Ile: Isoleucine; Leu: Leucine; Lys: Lysine; Met: Methionine; Phe: Phenylalanine; Thr: Threonine; Val: Valine; Arg: Arginine; Asp: Aspartate; Glu: Glutamate; Ser: Serine; Gly: Glycine; Ala: Alanine; Tyr: Tyrosine; Pro: Proline).

The amino acid composition of NF-HOB community biomass was compared to that of the NF-HOB community enriched by Hu et al. and soybean meal as reported in the literature (<u>Hu et al., 2020</u>; <u>Karr-Lilienthal et al., 2005</u>). Although the proportion of essential and non-essential amino acids in the high-titer biomass in this study was slightly lower, its composition closely resembled that of soybean meal, exhibiting higher levels of valine, glycine, and alanine (Fig. 3C).

### Appropriate setting of hydraulic retention time enables the feasibility of continuous cultivation

The specific growth rate of NF-HOB is evidently lower than that of ordinary HOB, making the previously commonly employed 10-hour HRT (<u>Matassa et al., 2016</u>) inapplicable. When HRT was set to 24 hours, only species with a specific growth rate of at least 0.042 were retained in the continuous reactor. The initial inoculation of biomass resulted in an initial COD_t_ of 186 mg/L; however, dilution over the next 4 days led to a daily reduction of half of the biomass in the reaction system. The average daily effluent COD_t_ (outlet) closely approximated the mean COD_t_ (inner) observed in the adjacent two-day system. Following re-inoculation to achieve an initial COD_t_ of 242 mg/L, a similar pattern of dilution was noted (Fig. 4A). Despite a decrease in the dilution rate, maintaining the inoculated biomass titer challenging when HRT was set to 36 hours. Furthermore, extending HRT to 48 hours resulted in NF-HOB accumulation. Once reaching a high density, it became difficult to reduce the HRT back to 36 hours while maintaining this density for more than a week; thus, 36 hours or less proved inadequate for supporting a stable reaction system. When HRT was set to 48 hours, the initial inoculation biomass measured COD_t_ at 99 mg/L. After a 2-day adaptation period, the community gradually enriched and achieved COD_t_ levels above 300 mg/L, reaching 328.58 mg/L on the 15th day. COD_s_ also accumulated progressively and accounted for over 25% of COD_t_ in later stages. The pH remained consistently stable within the range of 7.10 to 7.45 (Fig. 4B). Therefore, under suitable conditions, NF-HOB exhibited a specific growth rate greater than 0.021. After 16 days of continuous cultivation, COD_t_ reached 383.65 mg/L and CDW reached 218.58 mg. The biomass obtained exhibited a carbon content of 42.6% and a nitrogen content of 8.07%, corresponding to an MP content of 50.44%. By collecting the solution containing this biomass concentration every 48 hours as the product, the volumetric productivity for MP was achieved at a rate of 2.297 mg/(L·h), surpassing that attained in batch culture. Although this value is significantly lower than the previously reported MP productivity of the ammonium-assimilation HOB community (<u>Matassa et al.,</u> <u>2016</u>), its predominance relies on the stable biomass concentration. Given that the HRT of 48 hours has been demonstrated to support continuous cultivation of NF-HOB communities, opting for a larger inoculum size will inevitably augment stable biomass concentrations, promising high production efficiency in future large-scale practice.

**Fig. 4.**
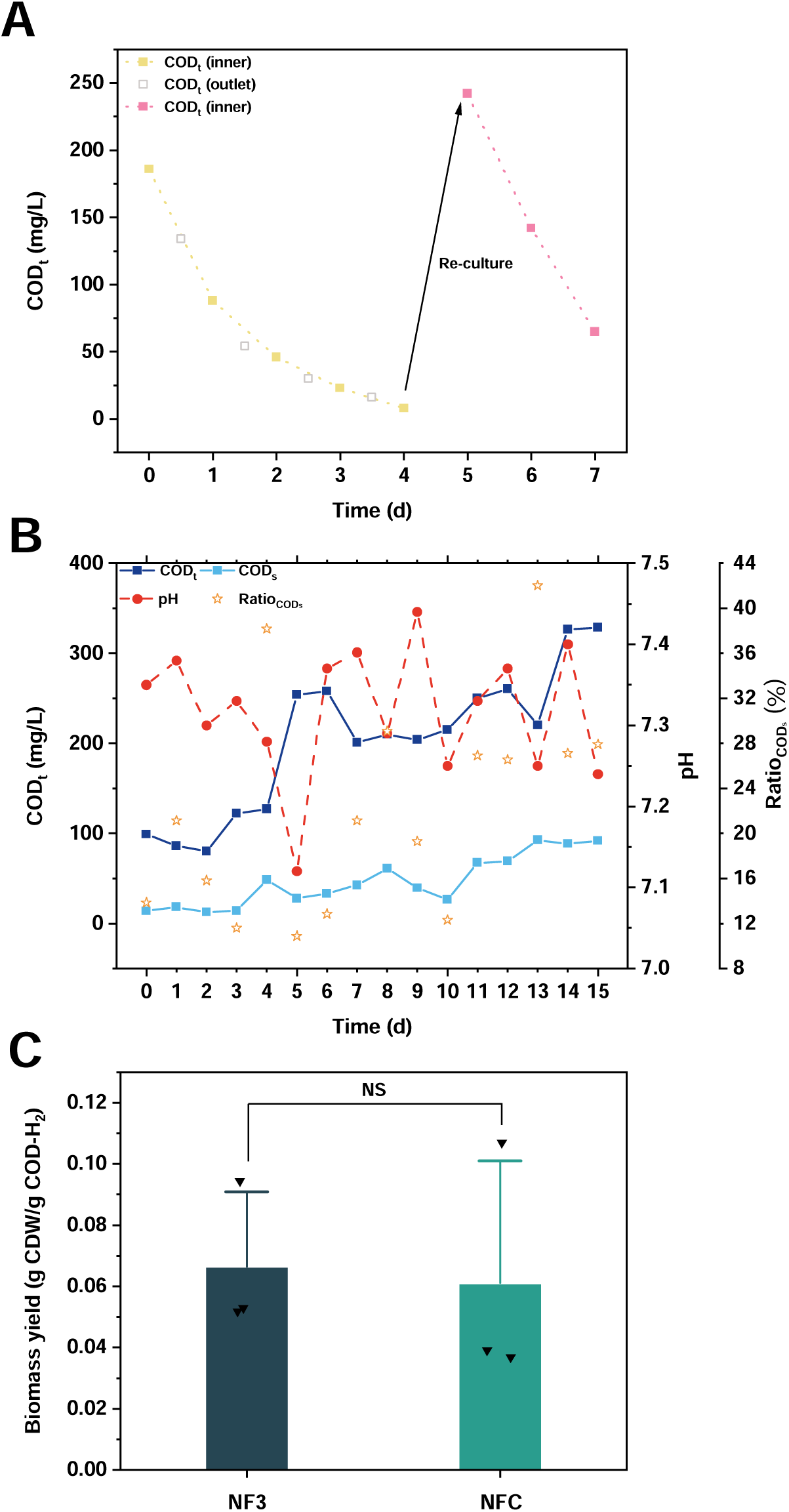
(A) COD_t_ concentration during continuous culture of the NF-HOB community at HRT = 24 h; COD_t_ (inner): COD_t_ of the liquid-phase reaction system; COD_t_ (outlet): daily average COD_t_ of liquid pumped out of the system. (B) Inner COD_t_ concentration, COD_s_ concentration, pH, and the ratio of COD_s_ to COD_t_ (Ratio_CODs_) during continuous culture of the NF-HOB community at HRT = 48 h. (C) Biomass yield of the NF-HOB community enriched during the third batch culture (NF3) and continuous culture (NFC) (Each value is the mean ± standard deviation of triplicates, “NS” indicates no significant difference because the two-tailed p-value of the t-test is > 0.05).

The headspace gas supply experiment conducted in sealed bottles (introduced in section 2.4) demonstrated that, although biomass growth of NF-HOB from continuous culture was comparatively lower within the same time frame (see supplementary material), the efficiency of biomass synthesis based on H_2_ was comparable to that obtained through sequencing batch culture (Fig. 4C). The range observed in this study (0.0369-0.1069 g CDW/g COD-H_2_) is consistent with previous findings.

### Microbial community analysis

The taxonomic composition of each enriched microbial community at the family and genus levels is illustrated in Fig. 5A and 5B, respectively. Among the community utilized as the inoculum source, the Xanthobacteraceae family constituted 76.7%, while *Xanthobacter* accounted for merely 2.70% and *Pseudoxanthobacter* accounted for only 1.32%. Consequently, 95% of Xanthobacteraceae species belonged to unidentified genera, classified as “others”.

**Fig. 5.**
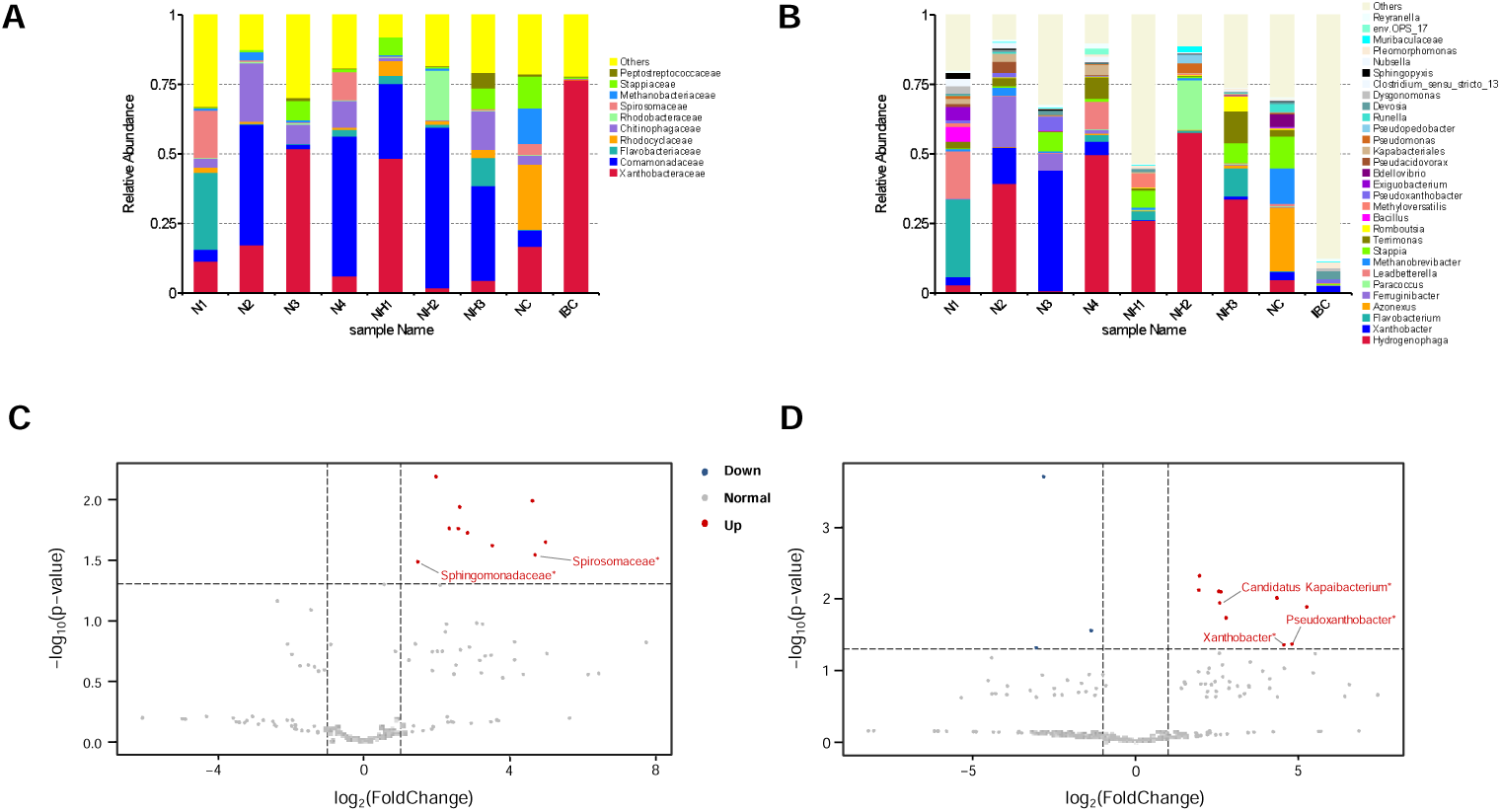
Microbial community structure and difference analysis: species distribution at the family (A) and genus (B) levels (IBC: HOB community used for sequencing batch culture experiments after initial domestication; N1: NF-HOB community enriched at R_H/O_ = 3.5; N2: NF-HOB community enriched during the first batch at R_H/O_ = 7; N3: NF-HOB community enriched during the second batch at R_H/O_ = 7; N4: NF-HOB community enriched during the third batch at R_H/O_ = 7; NH1: ammonium-assimilation HOB community enriched at R_H/O_ = 3.5; NH2: ammonium-assimilation HOB community enriched during the first batch at R_H/O_ = 7; NH3: ammonium-assimilation HOB community enriched during the second batch at R_H/O_ = 7; NC: NF-HOB community enriched during the continuous culture); volcano plot of species with significant abundance differences between NF-HOB and ammonium-assimilation HOB communities at the family (C) and genus (D) level (“Up” indicates that the abundance of the species represented by this point is significantly higher in the NF-HOB community, while “Down” indicates the opposite).

*Hydrogenophaga* has consistently maintained dominance as the genus in ammonium-assimilation HOB communities. Under low R_H/O_ conditions, Bacteroidota (53.6%) dominated the NF-HOB community, with *Flavobacterium* and *Leadbetterella* exhibiting relative abundances of 27.8% and 16.8%, respectively. *Flavobacterium* has been reported to possess capabilities for H_2_ oxidation (<u>Maimaiti et al., 2007</u>) and N_2_ fixation (<u>Kämpfer et al., 2015</u>; <u>Sorkhoh et al., 2010</u>); however, there is insufficient evidence to substantiate its CO_2_ fixation ability. Currently, *Leadbetterella* remains poorly understood. Therefore, the role of CO_2_ fixation may be assumed by *Bacillus* (<u>Maheshwari et al., 2017</u>), *Xanthobacter* (<u>Lu et al., 2020</u>), *Hydrogenophaga* (<u>Ehsani et</u> <u>al., 2019</u>; <u>Zhang et al., 2022</u>), etc.

Under high R_H/O_ conditions, the phylum Proteobacteria regained dominance. In the first batch, *Hydrogenophaga* (39.3%), *Ferruginibacter* (18.1%), and *Xanthobacter* (13.1%) were enriched. Although similar COD_t_ growth and ^Δ^TN_biomass_ were observed in the second and third batches, significant differences in community structure were noted. In the second batch, *Xanthobacter* emerged as the predominant genus, with a relative abundance of 43.3%. Simultaneously, *Pseudoxanthobacter*, also from the Xanthobacteraceae family, displayed a relative abundance of 4.9%. Moreover, *Stappia*, with a relative abundance of 6.8%, may occupy an ecological niche due to its potential for CO_2_ fixation (<u>Weber and King, 2007</u>; <u>Zhang et al., 2020a</u>); however, the precise role of *Ferruginibacter* (6.1%) remains unclear. In the third batch, *Hydrogenophaga* was re-enriched; additionally, *Leadbetterella*, *Terrimonas*, and *Xanthobacter* were identified as other genera with relative abundances exceeding 4%. Notably, the N_2_ fixation capability of *Terrimonas* has recently been emphasized (<u>Guo et al., 2024</u>), providing insights into its functional significance within the NF-HOB community.

The structure of the NF-HOB community obtained under continuous cultivation remained relatively complex, with genera exhibiting relative abundances exceeding 4%, including *Azonexus* (23.0%), *Methanobrevibacter* (12.7%, specifically *Methanobrevibacter arboriphilus*), *Stappia* (11.5%), *Hydrogenophaga* (4.9%), and *Bdellovibrio* (4.6%, predatory bacteria). However, it is noteworthy that apart from *Xanthobacter* with a relative abundance of 2.9% and a few species of *Bradyrhizobium* and *Pseudoxanthobacter*, there were numerous unidentified members belonging to the Xanthobacteraceae family, collectively accounting for over 13%. The *Azonexus* strain has been reported to possess both H_2_ oxidation and N_2_ fixation capabilities but lacks a CO_2_ fixation pathway (<u>Hu et al., 2022</u>). *Methanobrevibacter arboriphilus* represents a prototypical hydrogenotrophic methanogenic archaea (<u>Sharma et al., 2023</u>), with certain strains exhibiting potential for N_2_ fixation (<u>Poehlein et al., 2018</u>). *Hydrogenophaga* sp. NFH-34 harbors genes associated with N_2_ fixation but is unable to perform this function for growth maintenance under hydrogen autotrophic conditions (<u>Hu et al., 2022</u>). Consequently, similar to *Stappia*, there is insufficient direct evidence supporting its N_2_ fixation capability in the NF-HOB community. In general, besides the incompletely identified members of the Xanthobacteraceae family potentially involved in concurrent H_2_ metabolism, CO_2_ fixation, and N_2_ fixation, other abundant species are likely to assume specific responsibilities and engage in symbiotic cooperation.

The NF-HOB and ammonium-assimilation HOB communities exhibited significant abundance differences in 10 families and 28 genera. For NF-HOB, families with relatively higher relative abundance (averaging over 1%) included Spirosomaceae and Sphingomonadaceae (Fig. 5C), while the genera included *Xanthobacter*, *Pseudoxanthobacter*, and *Candidatus Kapaibacterium* (Fig. 5D). This observation can be attributed to the N_2_ fixation capability of *Xanthobacter* and *Pseudoxanthobacter* (<u>Arun et al., 2008</u>).

The abundance of *Hydrogenophaga* showed a significant positive correlation (p < 0.01) with the abundance of various functional enzymes, including three hydrogenases (EC:1.12.1.2, EC:1.12.7.2, EC:1.12.99.6), carbonic anhydrase, glutamate dehydrogenase (EC:1.4.1.3), glutamate synthase, and glutamine synthase (Fig. 6A). This underscores the pivotal role played by this genus in processes such as H_2_ oxidation and amino acid synthesis in NF-HOB communities. Among the other genera within the top 30 relative abundances, *Pseudopedobacter* and *Nubsella* displayed significant positive correlations (p < 0.05) with nitrogenase abundance. Additionally, *Nubsella* also demonstrated a strong correlation (p < 0.01) with hydrogenase (EC:1.12.1.2) abundance, as well as glutamate dehydrogenase (EC:1.4.1.3) and glutamate synthase. The abundance of *Muribaculum* showed a significantly positive association with Rubisco (p < 0.05), but the underlying mechanism is unknown.

**Fig. 6.**
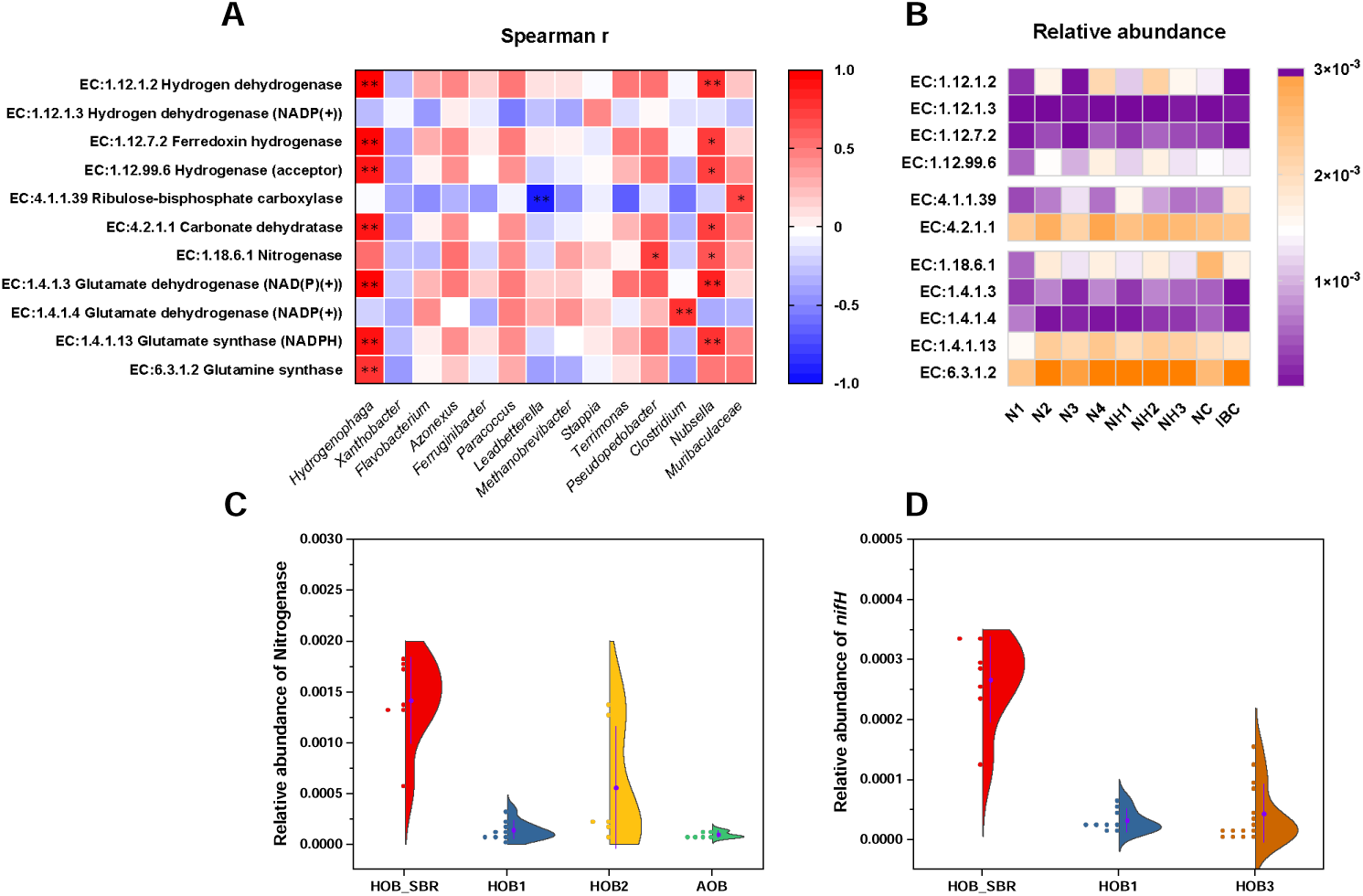
(A) Heatmap of Spearman correlation coefficients between the abundance of key functional enzymes and the abundance of dominant genera (* indicates the two-tailed p-value of the t-test is < 0.05, ** indicates the two-tailed p-value of the t-test is < 0.01). (B) Relative abundance of functional enzymes. (C) Comparison of the relative abundance of nitrogenase in microbial communities (HOB_SBR: HOB communities enriched during sequencing batch culture in this study; HOB1: previously reported NF-HOB communities; HOB2: previously reported N_2_-fixing bacterial communities enriched in H_2_-producing electrolytic cells; AOB: previously reported ammonium-oxidizing functional bacterial communities). (D) Comparison of the relative abundance of *nifH* in microbial communities (HOB3: ammonium-assimilation HOB communities). (In the violin plot, the purple dots indicate the mean value, and the length of the whisker lines represents the size of the standard deviation.)

The relative abundance of key functional enzymes in each HOB community is illustrated in Fig. 6B. The impact of nitrogen source supply conditions on these abundances was not decisive, which aligns with previous research findings (Wang et al., 2024). Notably, the NF-HOB community enriched through continuous culture exhibited a remarkably high relative abundance of nitrogenase at 0.26%, surpassing that obtained through sequencing batch culture by at least 0.45 times. This phenomenon may be crucial for the community to adapt to the prescribed dilution rate and thrive, leading to the proliferation of distinct dominant genera. Moreover, even the abundance of nitrogenase obtained through sequencing batch culture offers significant advantages over previously investigated NF-HOB communities (HOB1) (<u>Wang et al.,</u> <u>2024</u>), as well as the N_2_-fixing communities enriched in the microbial electrochemical system where water was electrolyzed for H_2_ production (HOB2) (<u>Zhang et al., 2022</u>; <u>Zhang et al., 2024</u>) and the ammonium oxidation functional bacterial communities (AOB) (<u>Fan et al., 2025</u>) (Fig. 6C). Similarly, the relative abundance of the representative gene encoding nitrogenase, *nifH*, surpassed that of both previous NF-HOB communities (HOB1) and ammonium-assimilation HOB communities (HOB3, for its community structure, see supplementary material) (Fig. 6D). This substantial nitrogenase abundance ensures efficient autotrophic N_2_ fixation to meet the demands for synthesizing MP.

### Outlook

Selecting a high R_H/O_ effectively enhanced the potential of NF-HOB, leading to remarkable achievements in MP synthesis during the later stage of sequencing batch culture, with a maximum titer reaching 983.26 mg/L, surpassing previous understanding. Other pivotal strategies implemented included ensuring a continuous supply of gas and optimizing the initial structure of the microbial community. Although the NF-HOB community exhibits a slightly lower proportion of MP and various essential amino acids in CDW compared to the *Xanthobacter* pure strain, the latter necessitates a headspace gas composition of 78% N_2_ and 2% O_2_ (<u>Hu et al., 2022</u>; <u>Sherbo et al., 2022</u>), which may present challenges for large-scale pure cultivation. This study validates that air can effectively serve as a source of both nitrogen and oxygen for promoting biomass growth, thereby bringing NF-HOB closer to practical application.

Continuous culture outperforms sequencing batch culture in terms of MP volumetric productivity or biomass yield for the ammonium-assimilation HOB community (<u>Matassa et al., 2016</u>). However, in this study, an advantage in volumetric productivity was found only for the NF-HOB community. To further enhance this production level (2.297 mg/[L·h]), several considerations should be addressed in future work. Firstly, the adoption of large-scale equipment with improved gas-liquid mass transfer efficiency is necessary. Prolonging the duration of continuous cultivation can enhance community stability and facilitate the identification of more NF-HOB strains. Research on the CO_2_ fixation performance of sulfur-oxidizing bacteria and nitrifying bacteria revealed that the extracellular free organic carbon (EFOC) inevitably produced exerted an inhibitory effect on the transcription of *cbb* genes, thereby impacting CO_2_ sequestration (<u>Zhang et al., 2021</u>; <u>Zhao et al., 2023</u>). This feedback mechanism should also be taken into account in the cultivation of NF-HOB, particularly given the significant proportion of observed COD_s_ under continuous cultivation mode. Development techniques for isolating EFOC may therefore be deemed a pivotal strategy. Additionally, the exploration of optimal parameters can be enhanced by introducing supplementary gradients within the HRT range of 36 to 48 hours.

## Conclusion

Under parallel operating conditions, a higher R_H/O_ benefited NF-HOB biomass growth, resulting in a comparable final titer to that of ammonium-assimilation HOB. For the NF-HOB community, a N_2_ fixation rate of 1.2-1.6 mg N_2_ per gram dry biomass per hour was observed in the last two batches, resulting in the highest MP titer of 983 mg/L, with an amino acid composition similar to that of soybean meal. Continuous culture confirmed the specific growth rate of NF-HOB ranged from 0.021 to 0.028, leading to enhanced MP volumetric productivity. Substantial nitrogenase abundance ensured efficient autotrophic N_2_ fixation to support MP synthesis.

### Supplementary material

E-supplementary data for this work can be found in e-version of this paper online.

---

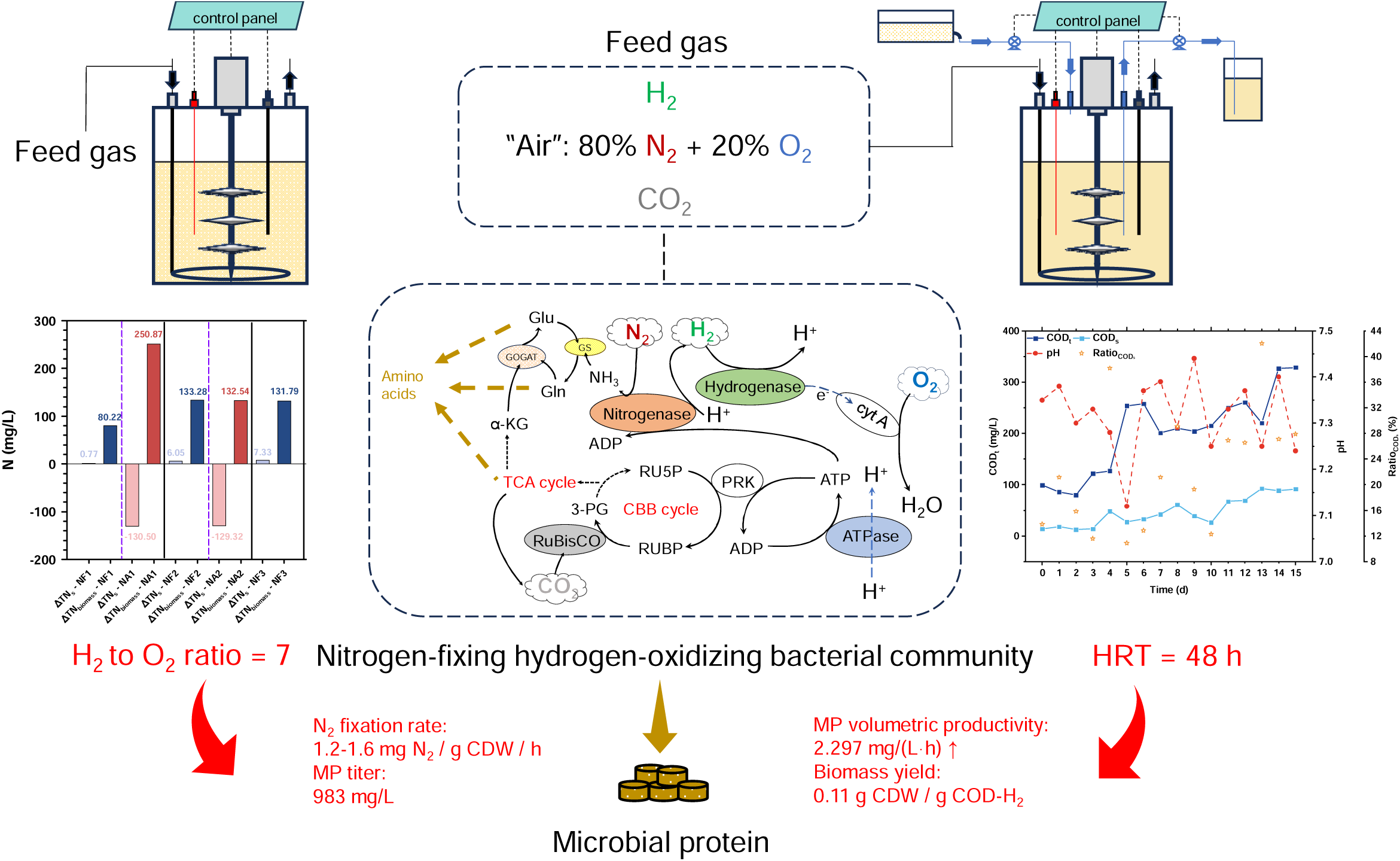

## Notes

https://www.researchgate.net/profile/Haoran-Wang-131/research

### Competing Interest Statement

The authors have declared no competing interest.

### Summary of Updates

Author affiliations updated; funding information updated; and also, when this article is submitted and accepted, there will be an updated list of authors.

### Funding

- the National Natural Science Foundation of China: 31970106
- the Sichuan Province Science and Technology Program: No. 2023SFSC1230

