## Supplementary material for "Enhancement strategies for the microbial protein production of nitrogen-fixing hydrogen-oxidizing bacterial community": 4 Supplementary Figures

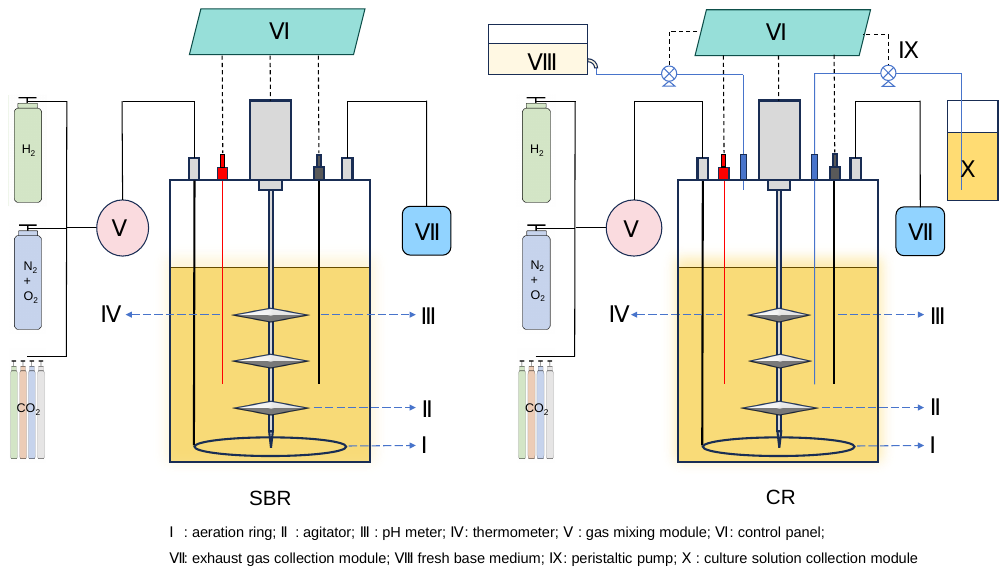

Fig. S1. Structural schematic diagram of sequencing batch reactor (SBR) and continuous reactor (CR).

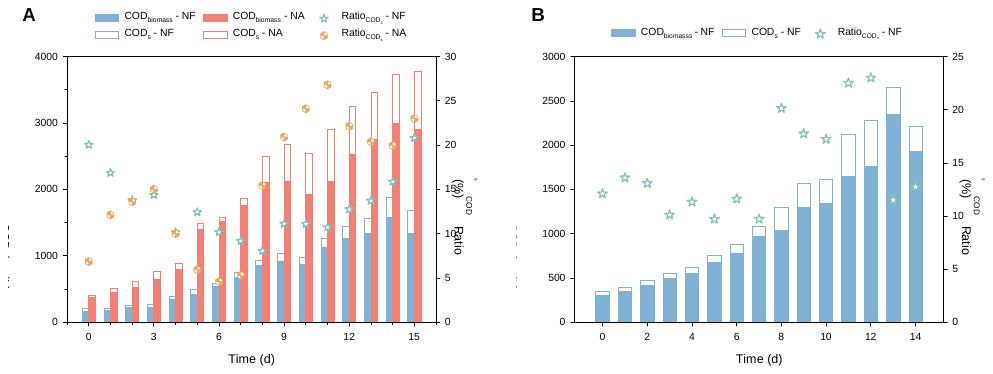

Fig. S2. Composition of COD_t_ (COD_s_ + COD_biomass_) in the second (A) and third (B) batches at R_H/O_ = 7 (Ratio_CODs_: the ratio of COD_s_ to COD_t_).

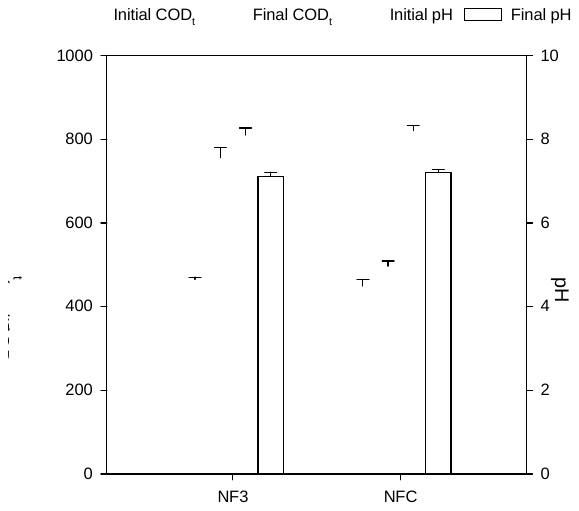

Fig. S3. Initial and final COD_t_ and pH of the headspace gas supply experiment to characterize gas consumption (NF3: the NF-HOB community enriched during the third batch culture; NFC: the NF-HOB community enriched during continuous culture; each value is the mean ± standard deviation of triplicates).

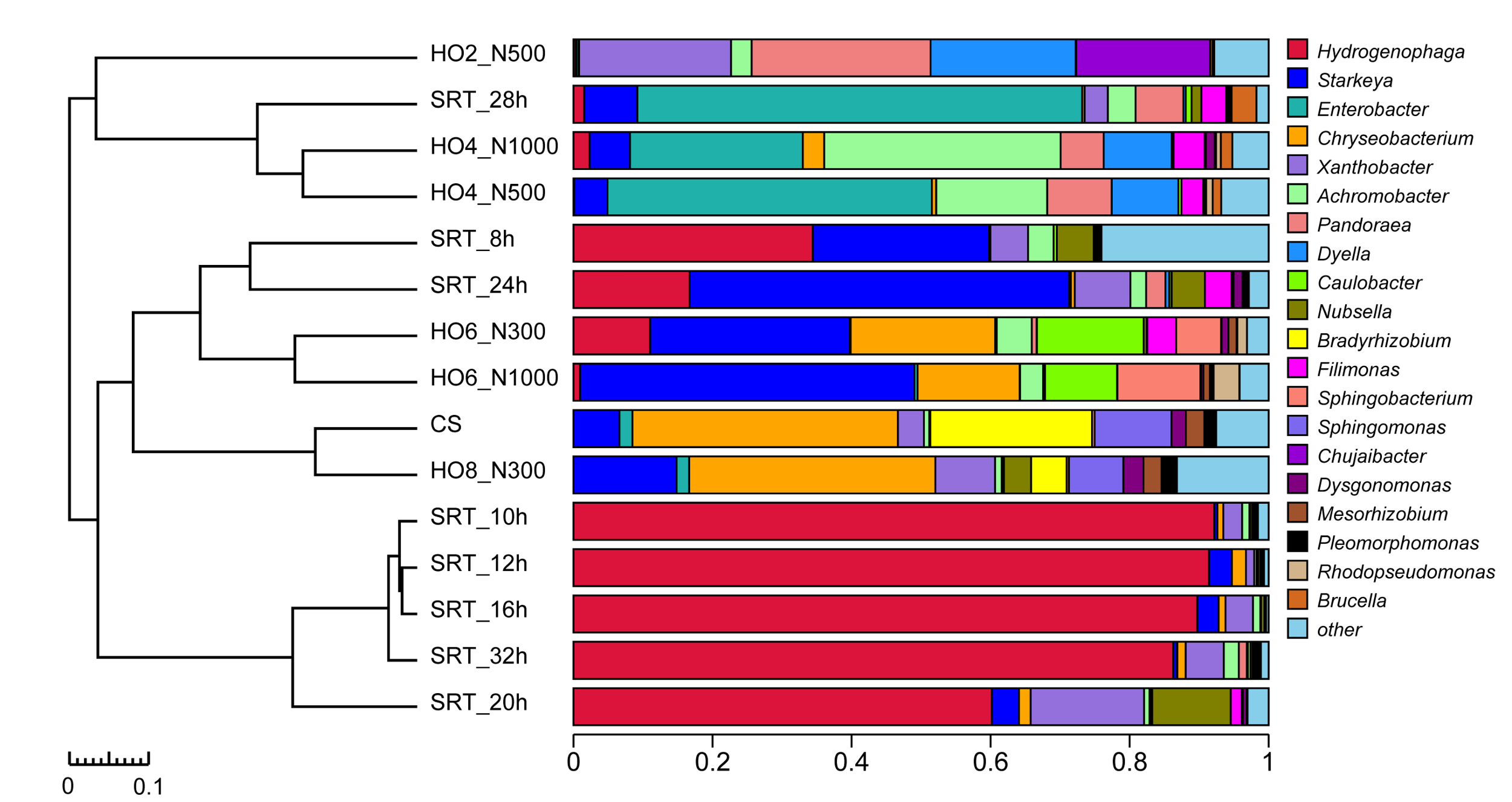
Fig. S4. Species distribution of "HOB3" communities (15 ammonium-assimilation HOB communities) at the genus level.
